# Context-Dependent Fitness Outcomes of Helping in the Cooperatively-Breeding Florida Scrub-Jay (*Aphelocoma coerulescens*)

**DOI:** 10.1101/2025.07.21.666021

**Authors:** Jeremy Summers, Bailey S.C.L. Jones, Elissa J. Cosgrove, Tori D. Bakley, Sahas Barve, Reed Bowman, John W. Fitzpatrick, Nancy Chen

## Abstract

Cooperatively breeding species frequently live in family groups of related individuals, with helpers delaying their own reproduction and participating in alloparental care, predator vigilance, and territory defense. It remains challenging to disentangle the roles of the indirect fitness benefits of helping kin and the potential direct fitness benefits helpers receive in the evolution of cooperative breeding. While many studies test for associations between helper relatedness and helping effort, few estimate the realized fitness consequences of helping in relation to these factors. Understanding these fitness outcomes elucidates the selective forces that maintain helping behavior, whether through inclusive fitness gains by helping related individuals or as a means of gaining access to later direct fitness benefits. Using 29 years of extensive demographic data from a closely monitored population of Florida scrub-jays (*Aphelocoma coerulescens*), we quantified the effect of helpers on breeder survival, offspring survival, and nestling production and how these effects depend on relatedness and sex of helpers. We found that female breeder survival was significantly greater when more helpers were present and that offspring survival was greater when more male helpers were present on small territories. Neither effect of helpers depended on the relatedness between helpers and the individuals they helped. Our results suggest that helping behavior is highly context-dependent and varies based on the potential impact of helping and the direct fitness benefits helpers receive.

## Introduction

In cooperatively breeding species, nonbreeding helpers delay reproduction and help raise offspring that are not their direct descendants (Hamilton 1964; Emlen 1982; Cockburn 1998; Hatchwell 1999). Helpers may contribute to activities such as incubation, food provisioning, predator vigilance, and territory defense, all of which can lead to increased survival and/or reproductive success of the dominant breeders (Crick 1992; Hatchwell 1999; Russell et al. 2007; Downing et al. 2020; Cardas et al. 2023). Given the potential fitness costs of delayed reproduction and helping behavior, multiple explanations for the evolution of cooperative breeding have been proposed (Cockburn 1998). Helping behavior may arise due to indirect fitness benefits accrued from helping relatives (kin selection) (Hamilton 1964; Russell and Hatchwell 2001; Griffin and West 2003; Hatchwell 2009). Alternatively, helping may provide direct fitness benefits from living in a larger group (group augmentation), increased future breeding opportunities, or experience gained via helping (skill hypothesis) (Selander 1965; Woolfenden 1975; Rood 1978; Magrath and Whittingham 1997; Kokko et al. 2001). The relative importance of indirect and direct benefits in the evolution of cooperative breeding remains controversial (Ligon and Stacey 1989; Cockburn 1998; Green et al. 2016; Downing et al. 2020; García-Ruiz et al. 2022).

Not all helpers contribute equally, as helping behavior is often biased by the perceived relatedness of helpers to the individuals they help (Komdeur 1994) and/or helper sex (Downing et al. 2018). Biases in helping behavior based on relatedness align with kin selection theory but could also result from other factors. Participation in helping behavior may be a means of signaling the quality of an individual to potential mates (social prestige hypothesis) (Zahavi 1995) or a requirement of remaining on the territory of dominant breeders (pay-to-stay hypothesis) (Gaston 1978; Kokko et al. 2001; Trapote et al. 2021). Sex differences in helping behavior also cannot be fully explained by kin selection because males and females are equally related to siblings they may help rear. Instead, sex-biased helping is thought to arise from sex differences in direct fitness, including differential likelihood of territory inheritance (Woolfenden and Fitzpatrick 1984; Downing et al. 2018).

Hypotheses explaining variation in helping effort are only cogent under the assumption that differences in helper effort lead to measurable variation in the fitness outcomes for the individuals being helped. However, quantifying the impact of helping contingent on sex and relatedness requires highly detailed demographic, relatedness, and downstream fitness data for both helpers and the individuals they help. These requirements have limited the ability for studies to investigate how variation among helpers affects helper impacts (but see Paquet et al. 2015, Koenig and Dickinson 2016, Leedale et al. 2018, Li et al. 2022, and Kerr et al. 2023). More studies combining helper number, relatedness, sex, and fitness outcomes are needed to elucidate how the direct and indirect fitness impacts of helping contribute to the evolution of cooperative breeding.

Here, we quantify how helper relatedness and sex affect breeder survival and reproductive success in the cooperatively breeding Florida scrub-jay (*Aphelocoma coerulescens*). Florida scrub-jays typically delay reproduction and help the socially dominant breeders rear offspring (Woolfenden and Fitzpatrick 1984). In addition to provisioning offspring (Stallcup and Woolfenden 1978; McGowan and Woolfenden 1990; Mumme 1992), helpers can contribute to sentinel behavior (McGowan and Woolfenden 1989; Hailman et al. 1994), predator mobbing (Francis et al. 1989), and territory defense (Woolfenden 1975). While most helpers remain on their natal territory, a few individuals disperse and act as helpers on other territories (Suh et al. 2022), resulting in mixed-kin family groups. Previous work has found positive associations between the presence or number of helpers and nest survival and fledgling production (Breininger et al. 2023), breeder survival (Woolfenden and Fitzpatrick 1984; Breininger et al. 2022), and post-fledging survival (Mumme et al. 2015).

Despite being a well-studied model of cooperative breeding, little is known about differences in the impacts of helpers based on relatedness or sex in the Florida scrub-jay. Florida scrub-jays exhibit sex-biased dispersal behavior and routes to breeding: females disperse farther and earlier than males (Aguillon et al. 2017; Suh et al. 2020), and males are more likely to inherit their natal territory (Woolfenden and Fitzpatrick 1984). While male Florida scrub-jay helpers display more helping effort (provisioning, sentinel behavior, and territory defense) (Stallcup and Woolfenden 1978; Hailman et al. 1994; Cardas et al. 2023), a previous study on offspring survival and reproductive success did not find a significant effect of helper sex (Mumme et al. 2015). Similarly, though there is documentation that Florida scrub-jay helpers prefer provisioning related nestlings (Mumme 1992), no study has investigated whether helper relatedness affects fitness outcomes (Kerr et al. 2023).

Here, we leverage an extensive, 32-year dataset from a long-term demographic study of Florida scrub-jays to test for differences in the effect of helpers on the fitness components that collectively result in lifetime reproductive success (breeder survival, nestling production, and offspring survival). We account for potential confounding variables such as breeder and territory quality, nest timing, and breeder age. We model both linear and non-linear effects of the number of helpers and explicitly test for effects of helper relatedness or sex on each fitness metric. Our study directly quantifies variation in the fitness effects of helping, allowing the evaluation of different models of cooperative breeding.

## Methods and Materials

### Study species and system

Florida scrub-jays are largely sedentary cooperative breeders restricted to the fire-maintained oak scrub of Florida (Woolfenden and Fitzpatrick 1984). They are socially monogamous, with one resident breeding pair occupying each territory. Offspring typically fledge 15-20 days after hatching, achieve sustained flight at ∼30 days, and reach nutritional independence at ∼90 days. Jays become physiologically capable of breeding at ∼300 days (yearling stage) and rarely disperse before they reach one year of age (Woolfenden and Fitzpatrick 1984; Mumme et al. 2015). Non-breeding adults (helpers) often delay dispersal and reproduction but help the resident breeding pair raise offspring (Woolfenden and Fitzpatrick 1984), though they may also disperse and “stage” by helping at territories other than their natal territory (Breininger et al. 2006; Suh et al. 2020).

We studied a color-banded population of Florida scrub-jays at Archbold Biological Station (27.10°N, 81.21°W) that has been continually and extensively monitored for five decades (see Woolfenden and Fitzpatrick, 1984, for detailed descriptions of the study area and population). Nestlings are banded on day 11 after hatching, and new immigrants are captured and banded shortly after they appear in the study tract. The population is censused monthly, allowing for long-term tracking of individuals from birth to death or disappearance from the study tract. All territories are mapped annually during the breeding season (March-June), and all nests are located and monitored until fledgling or failure (Woolfenden and Fitzpatrick 1984). Individual sex is assigned based on breeding behavior and sex-specific vocalizations or, starting in 1999, molecular sexing assays (Woolfenden and Fitzpatrick 1984; Chen et al. 2016). Florida scrub-jays have extremely low rates of extra-pair paternity (Quinn et al. 1999; Townsend et al. 2011), allowing us to generate a detailed and accurate pedigree from field observations. We confirmed pedigree relationships with genomic data in 1989-1991, 1995, and 1999-2013 (Chen et al. 2016).

We analyzed data from 1990-2021 to avoid biases caused by expansion in the study tract area before 1990. We excluded all years in which >10% of territories had unsexed helpers present during the breeding season (1994, 1997, and 1999) to avoid biases caused by missing sex data, and removed the remaining territory-years with unsexed helpers or where the nest was never found (*N* = 50). We additionally. For territory-years where breeders re-paired during the breeding season (*N* = 70), we selected the first breeding pair for our analyses, as the breeding status of the replacement breeder is dependent on the death or disappearance of a member of the first pair. We also excluded annual records of breeders who were censused on multiple territories during the breeding season (*N* = 46) to ensure we accurately matched breeders with helper counts. For our offspring survival and nestling production analyses, we excluded all territories-years where multiple broods of 11-day-old nestlings were hatched (*N* = 85), as second broods are typically attempted after the failure of the first brood (Woolfenden and Fitzpatrick 1984) and thus aren’t independent of the survival of the previous brood and would inflate nestling counts for pairs experiencing brood failure. Our final dataset included 1,782 unique territory-years, with 2,101 total nests.

### Fitness Measures

We considered three components of fitness: (1) breeder survival, (2) offspring survival, and (3) nestling production. We quantified annual breeder and offspring survival based on survival to the start of the breeding season (March) of the following year and modeled male and female breeder survival separately. We measured annual nestling production as the total number of 11-day-old nestlings produced by a pair during the breeding season.

### Helper Counts

We used detailed census data to count the number of helpers present on a territory for each fitness component of interest. All non-breeding adults (individuals >300 days old) observed on a territory were counted as helpers. For breeder survival, we counted all helpers present on the territory during the first month a breeder was observed for a given breeding season (March for most breeders). For offspring survival, we counted all helpers present on the territory the month the offspring hatched. For nestling production, we counted all helpers present on the territory during the first month the breeding pair was observed that breeding season.

### Statistical Analyses

We used generalized linear mixed models (GLMMs) to quantify the effect of helpers on breeder survival, offspring survival, and nestling production. For each fitness metric, we fitted a base model with a fixed effect of the total number of helpers followed by three additional models to investigate (1) any non-linear effect of helpers, such as diminishing returns of helping, (2) the importance of helper sex, and (3) the importance of helper relatedness to the breeders and nestlings they help. We set our helper sex and relatedness terms as deviations from the effect size estimate for the number of helper term based on the ratio of female helpers to total helpers or related helpers to total helpers, respectively. We calculated relatedness coefficients from the population pedigree using the ‘kinship’ function in the kinship2 package (Sinnwell et al. 2022). We categorized helpers as related if the relatedness coefficient between the helper and the focal breeder or offspring was ≥ 0.25, as the vast majority (94.5%) of related helpers were first or second-degree relatives to the individuals they helped.

To account for other factors that are known to influence survival and reproduction in our population, we included several territory- and nest-specific fixed effects in our models. All models included territory size, as larger territories have been shown to correlate with increased reproductive output (Woolfenden and Fitzpatrick 1984). For breeder survival, we also included breeder age as a fixed effect because this species exhibits actuarial senescence (McDonald et al. 1996). We added a quadratic term for breeder age to account for the non-linear relationship between age and mortality caused by lower survival of the youngest breeders (Woolfenden and Fitzpatrick 1984). For offspring survival and nestling production, we included breeding pair experience because reproductive success increases with greater pair experience in the Florida scrub-jay: new pairs with helpers produced half as many nestlings as experienced pairs with helpers (Woolfenden 1975). We also included hatch date for offspring survival and the earliest observation of a breeding pair for nestling production to control for seasonal declines in nest success (Woolfenden and Fitzpatrick 1984; Bowman and Woolfenden 2001). Finally, to account for inbreeding depression (Chen et al. 2016), we included individual inbreeding coefficients calculated from the population pedigree with the ‘calcInbreeding’ function in the pedigree package (Coster 2022) in the breeder and offspring survival models and pairwise relatedness coefficients between mates calculated with the ‘kinship’ function in the kinship2 package (Sinnwell et al. 2022) in the nestling production models.

We modeled breeder and offspring survival with binomial distributions. Breeder survival models included random effects for year, territory, and individual identity.

Offspring survival models included random effects for year, territory, and nest identity. We modeled nestling production using a zero-inflated generalized Poisson distribution after a simulation test for zero-inflation and a subsequent dispersion test indicated both significant zero-inflation (ratio of observed/simulated zeros = 1.73, *p* < 0.0001) and underdispersion (dispersion = 0.69, *p* < 0.0001) in our base model. Nestling production models included random effects of year, territory, and parent pair identity. Sample sizes for each model are included in Supplementary Tables S1-S7.

We checked for multicollinearity using variance inflation factors using the ‘vifstep’ function from the usdm package (Naimi et al. 2014). All models were fitted using the glmmTMB package (Brooks et al. 2017). While glmmTMB typically uses a maximum likelihood estimation method, priors can be set using a maximum *a priori* estimate (Brooks et al. 2017). Due to singularities in the random effect estimates for the female breeder survival models, we set a half-normal prior distribution for random effect estimation for all breeder survival models. We tested the uniformity and dispersion of our model residuals and tested for outliers using the ‘testResiduals’ function from the DHARMa package (Hartig 2024). We compared model performance using Akaike information criteria (AIC; Akaike 1998), and designated models with ΔAIC < 2 as performing equivalently. Model predictions were generated using the ‘ggpredict’ function from the ggeffect package (Lüdecke 2018), with bias correction for back-transforming whole-sample-level predictions. We conducted all statistical analyses in R version 4.5.0 (R Core Team 2025).

## Results

### Variation in Helper Presence, Sex Ratio, and Relatedness

The number of helpers on a territory varied depending on both breeder age and territory size. The total number of helpers ranged between 0-6, with helpers present on 52.3% of territories. The average number of helpers was 1.84, and helper number was positively correlated with breeder age (Spearman’s ρ = 0.23, *p* < 0.0001) and territory size (Spearman’s ρ = 0.19, *p* < 0.0001). Among nests with banded nestlings (11-day-old offspring), the number of helpers was negatively correlated with hatch date (Spearman’s ρ = -0.132, *p* < 0.0001). Overall, we found a similar number of female and male helpers, with an average sex ratio of 50:50, although larger territories had greater proportions of female helpers (Spearman’s ρ = 0.066, *p* = 0.0098). The majority of helpers were closely related (relatedness coefficient >= 0.25) to the breeders (68%) and offspring (84%) they helped. Breeder age was positively correlated with the proportion of related helpers (Spearman’s ρ = 0.44, *p* < 0.0001), and male breeders were more related to their helpers than female breeders (proportion of related helpers = 0.7 (male), 0.62 (female); Mann-Whitney-Wilcoxon W = 273546, *p* = 0.0005). Distributions of helper counts based on helper sex and relatedness can be found in Supplementary Figure S1.

### Helper Contributions to Breeder Survival

All models for both female and male breeder survival performed equally well (ΔAIC < 2). The simplest model for each sex showed that the survival of female breeders but not male breeders linearly increased with the total number of helpers (Fig. 1, Supp. Table S1-2). We found no evidence for a non-linear effect of helper number or any sex- or relatedness-dependent effects of helping on breeder survival.

**Figure 1.**
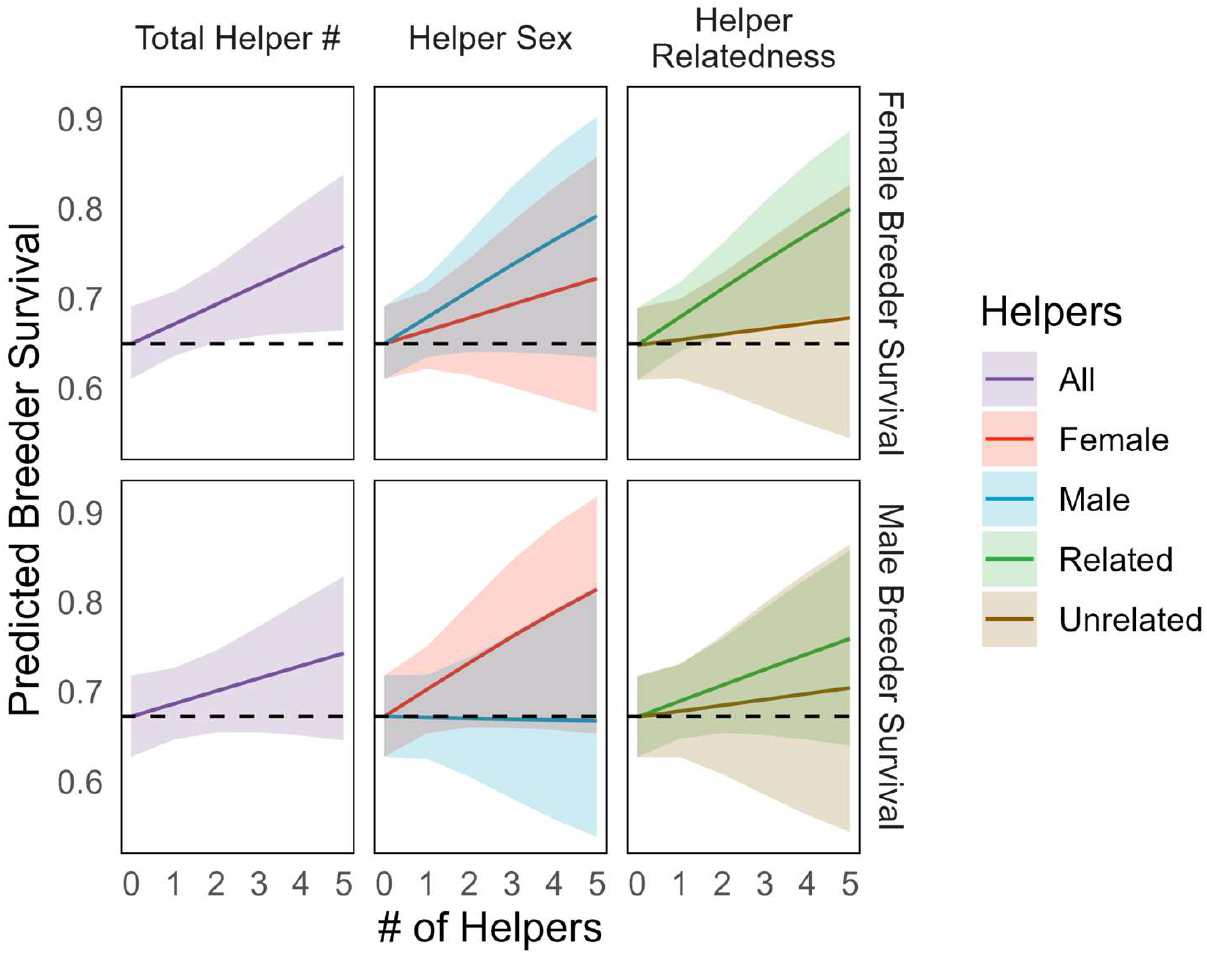
Predicted effect of the number of total (purple), female (red) and male (blue), or related (green) and unrelated (brown) helpers on annual survival of female (top) and male (bottom) breeders. Helpers are considered related if the relatedness coefficient between the helper and the focal breeder ≥ 0.25. Black horizontal dashed lines indicate the predicted survival probability with no helpers present, and shaded areas depict 95% confidence intervals. Helpers significantly increase female breeder survival.

### Helper Contributions to Offspring Survival

All models for offspring survival performed equally well (ΔAIC < 2), with the simplest model showing a positive linear effect of the total number of helpers on annual offspring survival (Fig. 2A, Supp. Table S3). As territory quality may affect helper contributions to offspring survival (Mumme et al. 2015), we further investigated the interaction between territory area and number of helpers. The best-performing model (ΔAIC = -2.5 relative to base model, Supp. Table S3-4) found that the effects of helpers on offspring survival depended on both territory size and helper sex, with a significant negative interaction between helper number and territory size and a positive interaction between helper sex ratio and territory size (Fig. 2B, Supp. Table S4). On small territories, offspring survival increased significantly with the number of male helpers, but on large territories, offspring survival was more influenced by the number of female helpers (Fig. 2B). Offspring survival continuously increased with territory size up to ∼20ha (Supp. Fig. S2) indicating that the presence of helpers may compensate for lower survival on smaller territories. Overall, our results indicate the presence of sex differences in helping, with male helpers having a greater effect on offspring survival in small territories and female helpers becoming more important in large territories. We found no support for non-linear helper effects on offspring survival, nor did helper effects vary based on helper relatedness.

**Figure 2.**
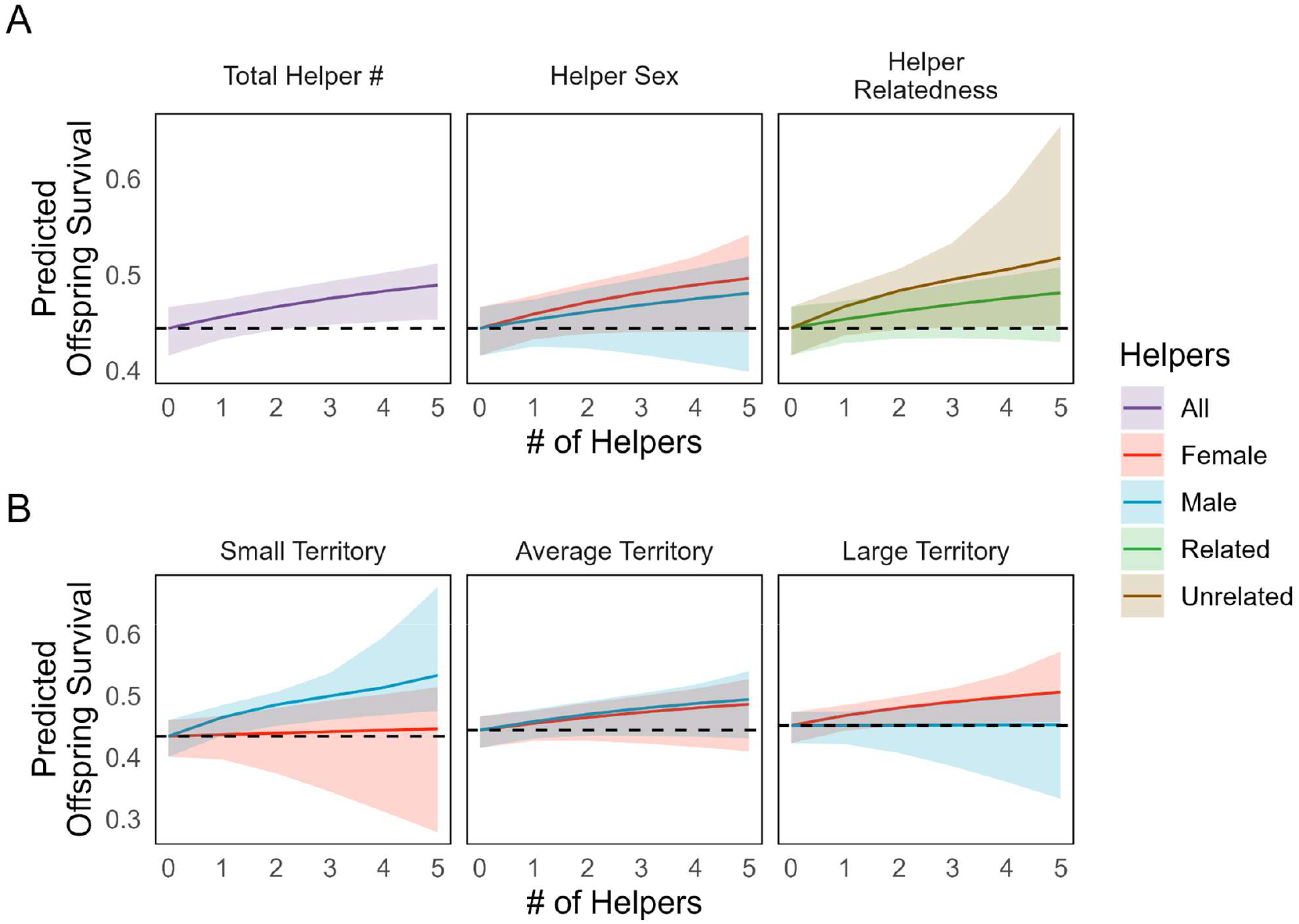
Predicted effect of the number of total (purple), female (red), and male (blue) helpers on offspring survival (A) with (red and blue) and without (purple) considering effects of helper sex and relatedness, and (B) with effects of helper sex and territory size. Helpers are considered related if the relatedness coefficient between the helper and the focal offspring ≥ 0.25. We show predictions for three territory sizes: small (first quantile, 9.6 ha), average (13.7 ha), and large (third quantile, 17.2 ha). Black horizontal dashed lines indicate the predicted survival probability with no helpers present, and shaded areas depict 95% confidence intervals. When ignoring territory size, the presence of helpers increases offspring survival, especially if helpers are unrelated. Offspring on small territories have greater survival when more male helpers are present, while offspring on large territories have greater survival when more female helpers are present.

### Helper Contributions to Nestling Production

The best-performing model for nestling production found a significant non-linear effect of the total number of helpers (Supp. Table S5). However, predicting the marginal impact of helper number revealed that nestling production does not significantly change with the number of helpers present (Supp. Fig. S3), suggesting that helpers do not contribute to nestling production.

## Discussion

Here, we investigate variation in helper contributions to breeder survival and reproductive success based on helper sex and relatedness in the Florida scrub-jay. We show that helpers increased annual survival of female breeders and of offspring, with helper contributions to offspring survival dependent on territory size and the sex of the helpers. Contrary to predictions of Hamilton’s rule (Hamilton 1964), we found no evidence that related helpers provided greater fitness benefits than unrelated helpers. Instead, we found evidence that direct fitness benefits may influence helping behavior: male helpers had a greater effect on offspring survival than female helpers, although this sex difference was only found on small territories. These results add to the growing literature on the context-dependent effects of helping behavior (Lejeune et al. 2016; Cousseau et al. 2022; García-Ruiz et al. 2022; Riehl and Smart 2022; Kerr et al. 2023) and demonstrate how multiple factors affect the fitness outcomes of helping.

Our results help reconcile conflicting estimates of the effect of helpers from previous work on the Florida scrub-jay. While some previous studies found that helpers improved breeder and offspring survival both in our study population at Archbold Biological Station (Woolfenden and Fitzpatrick 1984; Fitzpatrick and Bowman 2016) and other populations (Breininger et al. 2022; Breininger et al. 2023), other studies found that helpers provided no fitness benefits (Breininger 1999), only limited fitness benefits (namely, helpers reduced nest predation but did not affect breeder survival, egg production, or hatching success; (Mumme 1992), or even negative effects on juvenile body mass on small territories (Mumme et al. 2015). These discrepancies in the literature can be explained by two factors: first, the effects of helping behavior vary according to which individuals (males or females) are providing and receiving help. Second, the fitness benefits of helpers can be masked by variation in territory quality, emphasizing the need for standardized methods for comparing helper effects across studies (Downing et al. 2020).

Our study is the first to document a positive effect of helpers on female breeder survival in the Florida-scrub jay. While the estimated effect size of helpers was similar across our male and female breeder survival models, the association was not significant in males due to greater uncertainty (Fig. 1). Helpers may increase female but not male breeder survival because of sex differences in reproductive labor, mortality risks, or costs of helpers. While the benefits of helpers to breeder survival are typically the same between sexes in cooperatively breeding birds (Downing et al. 2020), our results match patterns found in the superb fairy wren (*Malurus cyaneus*) and sociable weaver (*Philetairus socius*). The superb fairy wren exhibits female-specific load lightening (Cockburn et al. 2008), including changes in provisioning rates and egg size (Russell et al. 2007). In the sociable weaver, the cost of helpers to male survival is thought to arise from competition between male breeders and their male helpers either for future breeding opportunities or for resources generally (Paquet et al. 2015). Both load-lightening in females or higher costs to males could be happening in the Florida scrub-jay. Previous work has documented sex differences in load-lightening in the presence of helpers: While both male and female Florida scrub-jay breeders provision young, only female breeders significantly reduce provisioning rates on territories with helpers (McGowan and Woolfenden 1990; Cardas et al. 2023). In addition, helpers might reduce the costs of reproduction or risk of predation for female breeders more than male breeders. Female breeders are the only group members that incubate (Woolfenden and Fitzpatrick 1984), and reduced nest predation involving mortality of the breeder has been cited as a consistently detectable benefit of helpers (Mumme 1992; Hailman et al. 1994; Niederhauser and Bowman 2014). In other cooperatively breeding birds, female breeders can regulate egg size in response to the presence of helpers, reducing resources used to produce eggs (Dixit et al. 2017). Finally, male breeders exhibit most of their aggressive behaviors toward their male helpers (Woolfenden and Fitzpatrick 1977). Future work might investigate differences in egg size or female breeder incubation and brooding behavior in response to the presence of helpers to determine the relative importance of female load-lightening.

Our finding that relatedness does not affect helper contributions, despite the heavily kin-structured nature of Florida scrub-jay family groups, provides further support for hypotheses that cooperative breeding in the Florida scrub-jay is maintained principally via direct fitness benefits (Woolfenden and Fitzpatrick 1984; Fitzpatrick and Bowman 2016). Indeed, our result lines up with recent arguments that kin-structuring in cooperative breeders arises from direct rather than indirect fitness. Rodrigues & Riehl (2025) and García-Ruiz et al. (2022) demonstrated that helping behavior can arise when helpers have no effect, or even negative effects, on breeder reproductive success. Limitations on breeding opportunities alone can result in sufficient selection against kin competition for breeders to tolerate mature offspring on their territories even if helpers provide no benefits (García-Ruiz et al. 2022; Rodrigues and Riehl 2025). Cooperatively breeding species with strict territorial limitations are more likely to help regardless of relatedness (Kingma 2017). The extreme habitat saturation at Archbold Biological Station (Woolfenden and Fitzpatrick 1984; Fitzpatrick and Bowman 2016) is likely to produce conditions that favor delayed dispersal and toleration of previous offspring and even non-kin helpers on the breeding territory. Our study is one of the few that can estimate how relatedness influences helping effects and directly evaluate the roles of direct and indirect fitness in the evolution of cooperative breeding (but see Dickinson et al., 1996, Green and Hatchwell, 2018, Hatchwell et al., 2004, and Kerr et al., 2023 for examples).

Sex differences in helping can arise from differences in the direct fitness benefits helpers receive. The “Philopatry hypothesis” (Capilla-Lasheras et al. 2024) predicts that the more philopatric sex is more likely to inherit its natal territory and should invest more in helping, ensuring access to the territory via a “pay-to-stay” system or improving future reproductive success through group augmentation (Kokko et al. 2001). In Florida scrub-jays, males are socially dominant over females and are more likely to inherit territory (Woolfenden and Fitzpatrick 1977; Woolfenden and Fitzpatrick 1984), leading to female-biased dispersal (Suh et al. 2020). We found that male helpers had greater effects on offspring survival, consistent with observations that male helpers provision at greater rates (Cardas et al. 2023).

We found that helper effects on offspring survival were dependent on territory size, with male helpers providing greater benefits on smaller territories (Fig. 2B). This result appears to contradict past research that suggested helpers compete with offspring for resources on smaller territories, reducing offspring body mass (Mumme et al. 2015). In our study, group size was positively correlated with territory size (Spearman’s ρ = 0.19, *p* < 0.0001), possibly reducing competition for resources on smaller territories. Our results may reflect the greater benefit that increased provisioning has on smaller territories, as provisioning rates are most likely to increase offspring survival when nestling starvation risks are high (Hatchwell 1999). In meerkats (*Suricata suricatta*), helping effort was more equal among helpers within smaller groups (Rotics and Clutton-Brock 2021), which may result in larger, and thus easier to detect, effects of individual helpers on offspring survival. Future work quantifying male helping effort across territory sizes is needed to determine whether the patterns we observe are caused by differences in the sensitivity of offspring survival to help or differences in helping behavior itself.

We note that our study focused on the fitness outcomes of helping, not variation in helping behavior itself. Future work can clarify how individual variation in helping behaviors contributes to the observed differences in fitness outcomes, which can help distinguish between the pay-to-stay and social prestige hypotheses (Kingma et al. 2011) and the group augmentation or inclusive fitness hypotheses (Hamilton 1964; Kokko et al. 2001). Furthermore, as helpers are more likely to disperse after the death of a related breeder (Suh et al. 2020), future work should jointly model breeder survival, helper survival, and helper dispersal to characterize how the dispersal of helpers, and thus their reproductive strategies, affects the individuals on their natal territory.

## Conclusion

Drawing upon 29 years of extensive demographic data from a population of Florida Scrub-Jays, we found that helpers increase female breeder and offspring survival, with no effect of genetic relatedness. The effect of helpers on offspring survival depended on complex interactions between helper sex and territory size, emphasizing how the fitness impacts of helpers differ depending on context. Our results suggest that differences in territory inheritance and dispersal behavior between males and females play a major role in shaping variation in helper contributions. These findings reinforce growing evidence of the context-dependent effects of helping behavior, where individual helpers balance the costs and benefits of helping within their environmental and social limitations, and provide additional evidence supporting the role of direct fitness benefits in the evolution of cooperative breeding (Kingma 2017; Downing et al. 2018; Rodrigues and Riehl 2025).

## Acknowledgments

We are grateful for all the interns and staff at Archbold Biological Station who collected demographic data on the Florida Scrub-Jay over the past half-century. We thank the members of the Chen Lab, especially Dr. Shailee Shah, and the Avian Ecology Program at Archbold Biological Station for their valuable feedback. BJ was supported by Swarthmore College’s David Baltimore/Broad Foundation Endowment, and JS was supported by NIH grant 1R35GM133412 to NC.

## Ethics statement

All fieldwork was approved by the Cornell University Institutional Animal Care and Use Committee (IACUC 2010-0015) and authorized by permits from the US Fish and Wildlife Service (TE8247238), the US Geological Survey (bandingpermit07732), and the Florida Fish and Wildlife Conservation Commission (LSSC-10-00205).

## Author contributions

Jeremy Summers: Conceptualization, Methodology, Software, Resources, Formal analysis, Investigation, Data Curation, Writing - Review and Editing.

Bailey S.C.L. Jones: Conceptualization, Methodology, Software, Formal analysis, Investigation, Data Curation, Writing - Original Draft, and Visualization.

Reed Bowman: Investigation and Data Curation. Sahas Barve: Investigation and Data Curation.

John W. Fitzpatrick: Investigation and Data Curation.

Nancy Chen: Conceptualization, Methodology, Investigation, Resources, Data Curation, Writing - Review and Editing, Supervision, Project Administration, and Funding Acquisition.

## Availability of data and materials

All data and code used in this study can be found at https://github.com/jtsummers53/FSJhelperEffect

