## Supplementary Information for "Context-Dependent Fitness Outcomes of Helping in the Cooperatively-Breeding Florida Scrub-Jay (*Aphelocoma coerulescens*)"

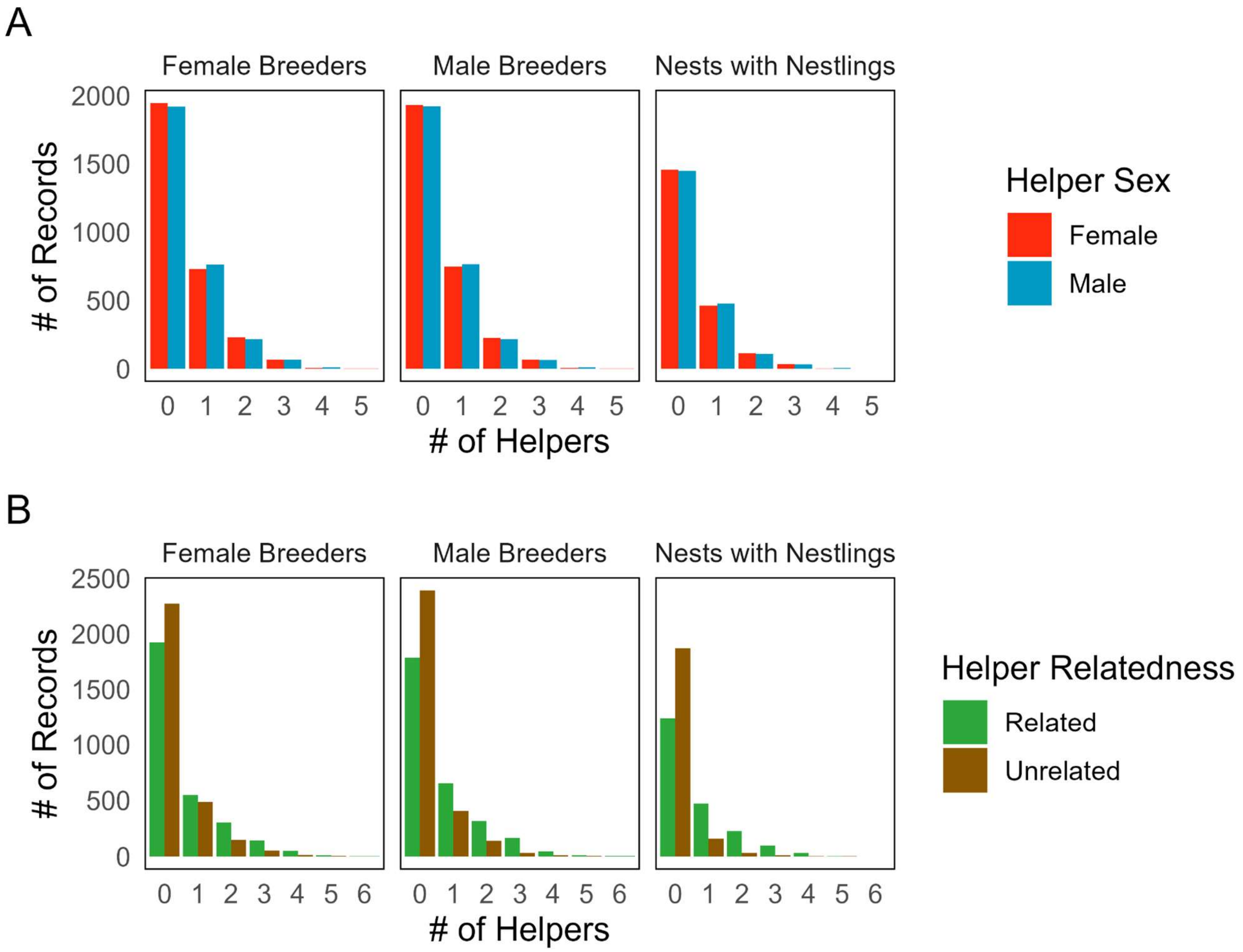


**Figure S1.** The number of helpers stratified by (A) sex and (B) genetic relatedness (relatedness coefficient ≥ 0.25) for female breeders, male breeders, and nests included in the offspring survival models. Helpers tend to be closely related to the focal individual.

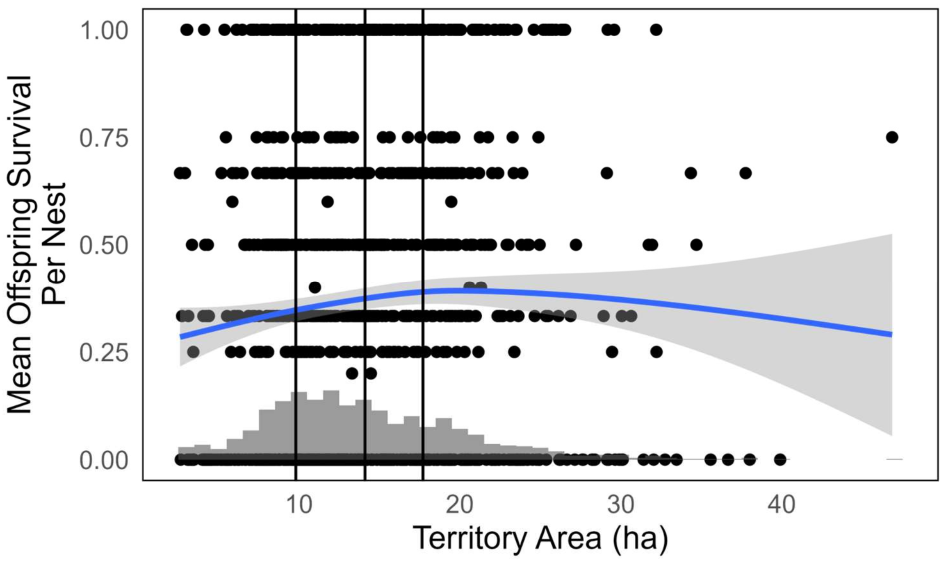


**Figure S2.** Offspring survival per nest varies with territory size. Black dots show data for individual nests, and the blue line is the weighted line-of-best-fit, and the gray histogram shows the distribution of territory sizes. Black vertical lines indicate the territory sizes used to generate predictions in Figure 2: the first quartile (9.8 ha), average (14.1 ha), and third quartile (17.7 ha). A higher proportion of offspring in a nest survive on larger territories up to ~20 ha in size.

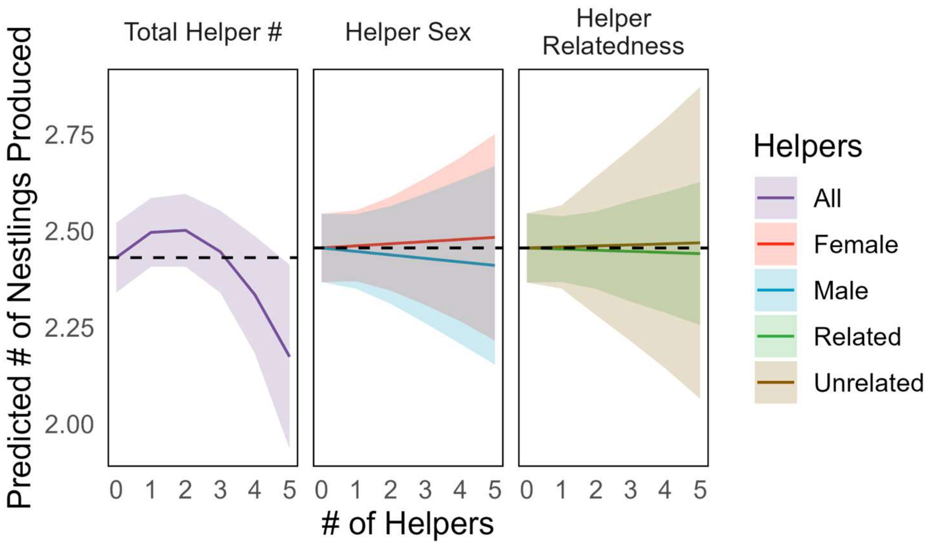


**Figure S3.** Predicted effect of the number of total (purple), female (red), and male (blue) helpers on nestling production (# of day 11 nestlings produced). Helpers are considered related if the relatedness coefficient between the helper and the focal breeder ≥ 0.25. Black horizontal dashed lines indicate the predicted # of day 11 offspring produced with no helpers present, and shaded areas depict 95% confidence intervals. Helpers did not significantly impact nestling production.

### Tables

| | Base | Non-Linear | Helper Sex | Helper Relatedness |
|---|---|---|---|---|
| Intercept | **0.686*** | **0.692*** | **0.689*** | **0.712*** |
| | (0.318) | (0.318) | (0.318) | (0.322) |
| Breeder Age | **0.290**** | **0.298**** | **0.290**** | **0.283**** |
| | (0.097) | (0.097) | (0.097) | (0.098) |
| Breeder Age$^2$ | **-0.029***** | **-0.030***** | **-0.029***** | **-0.029***** |
| | (0.007) | (0.007) | (0.007) | (0.007) |
| Inbreeding Coef. | -1.695 | -1.615 | -1.704 | -1.738 |
| | (3.787) | (3.786) | (3.778) | (3.778) |
| Territory Area | 0.012 | 0.013 | 0.012 | 0.012 |
| | (0.012) | (0.013) | (0.012) | (0.012) |
| # of Helpers | 0.081 | -0.066 | -0.005 | 0.037 |
| | (0.064) | (0.156) | (0.098) | (0.111) |
| # of Helpers$^2$ | | 0.045 | | |
| | | (0.044) | | |
| Helper Sex Ratio | | | 0.176 | |
| | | | (0.153) | |
| Related Helper Ratio | | | | 0.064 |
| | | | | (0.134) |
| Year | 0.253 | 0.253 | 0.258 | 0.254 |
| Territory ID | 0.335 | 0.335 | 0.342 | 0.333 |
| Individual ID | 0.506 | 0.504 | 0.494 | 0.500 |
| Num.Obs. | 1489 | 1489 | 1489 | 1489 |
| R2 Marg. | 0.047 | 0.049 | 0.048 | 0.047 |
| R2 Cond. | 0.158 | 0.159 | 0.158 | 0.156 |
| AIC | 1499.5 | 1500.4 | 1500.2 | 1501.3 |
| BIC | 1547.3 | 1553.5 | 1553.3 | 1554.4 |
| RMSE | 0.38 | 0.38 | 0.38 | 0.38 |

* $p < 0.05$, ** $p < 0.01$, *** $p < 0.001$

**Table S1**. Model results for male breeder annual survival. We fitted models that included (1) the total number of helpers (Base Model), (2) a quadratic term for total number of helpers (Non-Linear Model), (3) a sex-specific helper term (Helper Sex Model), and (4) a relatedness-specific helper term (Helper Relatedness Model). We present $\beta$ estimates for fixed effects with the standard error below in parentheses, standard deviations for random effects, sample sizes, and information on model fits.

| | Base | Non-Linear | Helper Sex | Helper Relatedness |
|---|---|---|---|---|
| Intercept | **1.496*** ** | **1.491*** ** | **1.492*** ** | **1.565*** ** |
| | (0.342) | (0.342) | (0.342) | (0.346) |
| Breeder Age | 0.062 | 0.059 | 0.063 | 0.039 |
| | (0.109) | (0.110) | (0.109) | (0.111) |
| Breeder Age$^2$ | -0.015 | -0.014 | -0.015 | -0.013 |
| | (0.008) | (0.008) | (0.008) | (0.008) |
| Inbreeding Coef. | -2.865 | -2.815 | -2.906 | -2.852 |
| | (4.585) | (4.587) | (4.587) | (4.566) |
| Territory Area | 0.001 | 0.000 | 0.001 | 0.001 |
| | (0.012) | (0.012) | (0.012) | (0.012) |
| # of Helpers | **0.125*** | 0.192 | 0.167 | 0.035 |
| | (0.063) | (0.147) | (0.099) | (0.097) |
| # of Helpers$^2$ | | -0.020 | | |
| | | (0.039) | | |
| Helper Sex Ratio | | | -0.084 | |
| | | | (0.151) | |
| Related Helper Ratio | | | | 0.144 |
| | | | | (0.121) |
| Year | 0.221 | 0.219 | 0.219 | 0.222 |
| Territory ID | 0.227 | 0.226 | 0.227 | 0.220 |
| Individual ID | 0.370 | 0.370 | 0.371 | 0.355 |
| Num.Obs. | 1489 | 1489 | 1489 | 1489 |
| R2 Marg. | 0.032 | 0.032 | 0.032 | 0.033 |
| R2 Cond. | 0.097 | 0.097 | 0.097 | 0.094 |
| AIC | 1484.8 | 1486.5 | 1486.5 | 1485.4 |
| BIC | 1532.5 | 1539.6 | 1539.5 | 1538.5 |
| RMSE | 0.38 | 0.38 | 0.38 | 0.38 |

* $p < 0.05$, ** $p < 0.01$, *** $p < 0.001$

**Table S2.** Model results for female breeder annual survival. We fitted models that included (1) the total number of helpers (Base Model), (2) a quadratic term for total number of helpers (Non-Linear Model), (3) a sex-specific helper term (Helper Sex Model), and (4) a relatedness-specific helper term (Helper Relatedness Model). We present $\beta$ estimates for fixed effects with the standard error below in parentheses, standard deviations for random effects, sample sizes, and information on model fits.

| | Base | Non-Linear | Helper Sex | Helper Relatedness |
|---|---|---|---|---|
| Intercept | 0.731 | 0.700 | 0.734 | 0.723 |
| | (0.426) | (0.427) | (0.426) | (0.426) |
| Hatch Day | **-0.015*** ** | **-0.015*** ** | **-0.015*** ** | **-0.015*** ** |
| | (0.003) | (0.003) | (0.003) | (0.003) |
| Pair Experience | 0.014 | 0.014 | 0.015 | 0.019 |
| | (0.031) | (0.031) | (0.031) | (0.031) |
| Inbreeding Coef. | -3.670 | -3.711 | -3.612 | -3.681 |
| | (2.251) | (2.255) | (2.254) | (2.255) |
| Territory Area | 0.012 | 0.011 | 0.012 | 0.012 |
| | (0.009) | (0.009) | (0.009) | (0.009) |
| # of Helpers | **0.113*** | 0.219 | 0.083 | **0.228*** |
| | (0.053) | (0.120) | (0.079) | (0.115) |
| # of Helpers$^2$ | | -0.034 | | |
| | | (0.035) | | |
| Helper Sex Ratio | | | 0.059 | |
| | | | (0.118) | |
| Related Helper Ratio | | | | -0.144 |
| | | | | (0.128) |
| Year | 0.455 | 0.455 | 0.456 | 0.455 |
| Territory ID | 0.333 | 0.331 | 0.333 | 0.344 |
| Nest ID | 0.745 | 0.745 | 0.745 | 0.741 |
| Num.Obs. | 2890 | 2890 | 2890 | 2890 |
| R2 Marg. | 0.024 | 0.024 | 0.024 | 0.024 |
| R2 Cond. | 0.228 | 0.229 | 0.229 | 0.229 |
| AIC | 3649.5 | 3650.6 | 3651.3 | 3650.3 |
| BIC | 3703.2 | 3710.2 | 3711.0 | 3709.9 |
| RMSE | 0.41 | 0.41 | 0.41 | 0.41 |

* $p < 0.05$, ** $p < 0.01$, *** $p < 0.001$

**Table S3.** Model results for offspring annual survival. We fitted models that included (1) the total number of helpers (Base Model), (2) a quadratic term for total number of helpers (Non-Linear Model), (3) a sex-specific helper term (Helper Sex Model), and (4) a relatedness-specific helper term (Helper Relatedness Model). We present $\beta$ estimates for fixed effects with the standard error below in parentheses, standard deviations for random effects, sample sizes, and information on model fits.

| | Base | Non-Linear | Helper Sex | Helper Relatedness |
|---|---|---|---|---|
| Intercept | 0.629 | 0.540 | 0.579 | 0.626 |
| | (0.436) | (0.439) | (0.436) | (0.436) |
| Hatch Day | **-0.015*** ** | **-0.015*** ** | **-0.015*** ** | **-0.015*** ** |
| | (0.003) | (0.003) | (0.003) | (0.003) |
| Pair Experience | 0.012 | 0.009 | 0.012 | 0.017 |
| | (0.031) | (0.031) | (0.031) | (0.031) |
| Inbreeding Coef. | -3.547 | -3.604 | -3.527 | -3.578 |
| | (2.255) | (2.251) | (2.261) | (2.261) |
| Territory Area | 0.019 | **0.023*** | 0.019 | 0.019 |
| | (0.011) | (0.012) | (0.011) | (0.011) |
| # of Helpers | 0.253 | **0.691*** | **0.645**** | 0.407 |
| | (0.139) | (0.316) | (0.221) | (0.346) |
| # of Helpers$^2$ | | -0.154 | | |
| | | (0.101) | | |
| Helper Sex Ratio | | | **-0.811*** | |
| | | | (0.351) | |
| Related Helper Ratio | | | | -0.200 |
| | | | | (0.380) |
| # of Helpers Territory Area Interaction | -0.009 | -0.033 | **-0.036**** | -0.012 |
| | (0.009) | (0.020) | (0.014) | (0.022) |
| # of Helper$^2$ Territory Area Interaction | | 0.008 | | |
| | | (0.007) | | |
| Helper Sex Ratio Territory Area Interaction | | | **0.055**** | |
| | | | (0.021) | |

| | | | | |
|---|---|---|---|---|
| Related Helper Ratio Territory Area Interaction | | | | 0.004 |
| | | | | (0.023) |
| Year | 0.454 | 0.448 | 0.460 | 0.454 |
| Territory ID | 0.335 | 0.339 | 0.330 | 0.346 |
| Nest ID | 0.744 | 0.740 | 0.737 | 0.741 |
| Num.Obs. | 2890 | 2890 | 2890 | 2890 |
| R2 Marg. | 0.024 | 0.026 | 0.029 | 0.025 |
| R2 Cond. | 0.229 | 0.228 | 0.230 | 0.230 |
| AIC | 3650.3 | 3651.7 | 3647.0 | 3653.2 |
| BIC | 3710.0 | 3723.4 | 3718.6 | 3724.8 |
| RMSE | 0.41 | 0.41 | 0.41 | 0.41 |

* p < 0.05, ** p < 0.01, *** p < 0.001

**Table S4.** Model results for offspring annual survival with interactions between helper number and territory size. We fitted models that included (1) the total number of helpers (Base Model), (2) a quadratic term for total number of helpers (Non-Linear Model), (3) a sex-specific helper term (Helper Sex Model), and (4) a relatedness-specific helper term (Helper Relatedness Model). We present $\beta$ estimates for fixed effects with the standard error below in parentheses, standard deviations for random effects, sample sizes, and information on model fits.

| Component | | Base | Non-Linear | Helper Sex | Helper Relatedness |
|---|---|---|---|---|---|
| Conditional | Intercept | **1.159*** ** | **1.150*** ** | **1.159*** ** | **1.159*** ** |
| | | (0.041) | (0.041) | (0.041) | (0.041) |
| | Breeding Start Month | **-0.138*** ** | **-0.136*** ** | **-0.138*** ** | **-0.138*** ** |
| | | (0.025) | (0.025) | (0.025) | (0.025) |
| | Pair Experience | **0.018** ** | **0.018** ** | **0.017** ** | **0.018** ** |
| | | (0.006) | (0.006) | (0.006) | (0.006) |
| | Breeder Relatedness | -0.639 | -0.677 | -0.633 | -0.638 |
| | | (0.418) | (0.417) | (0.417) | (0.418) |
| | Territory Area | 0.000 | -0.000 | 0.000 | 0.000 |
| | | (0.002) | (0.002) | (0.002) | (0.002) |
| | # of Helpers | -0.001 | 0.039 | -0.004 | 0.001 |
| | | (0.009) | (0.021) | (0.014) | (0.021) |
| | # of Helpers$^2$ | | **-0.012*** | | |
| | | | (0.006) | | |
| | Helper Sex Ratio | | | 0.006 | |
| | | | | (0.021) | |
| | Related Helper Ratio | | | | -0.002 |
| | | | | | (0.023) |
| | Year | 0.030 | 0.029 | 0.030 | 0.030 |
| | Territory ID | 0.050 | 0.050 | 0.050 | 0.050 |
| | Pair ID | 0.021 | 0.020 | 0.014 | 0.021 |
| Zero-Inflated | Intercept | **-0.931*** ** | **-0.931*** ** | **-0.931*** ** | **-0.931*** ** |
| | | (0.060) | (0.060) | (0.060) | (0.060) |
| Dispersion | | 0.294 | 0.292 | 0.295 | 0.294 |
| | Num.Obs. | 1428 | 1428 | 1428 | 1428 |

| | | | | |
|---|---|---|---|---|
| R2 Marg. | 0.013 | 0.013 | 0.012 | 0.013 |
| R2 Cond. | 0.020 | 0.021 | 0.019 | 0.020 |
| AIC | 4399.6 | 4397.6 | 4401.6 | 4401.6 |
| BIC | 4457.5 | 4460.7 | 4464.7 | 4464.8 |
| RMSE | 1.45 | 1.45 | 1.45 | 1.45 |

* p < 0.05, ** p < 0.01, *** p < 0.001

**Table S5.** Model results for nestling production. We fitted models that included (1) the total number of helpers (Base Model), (2) a quadratic term for total number of helpers (Non-Linear Model), (3) a sex-specific helper term (Helper Sex Model), and (4) a relatedness-specific helper term (Helper Relatedness Model). Model components are divided into Conditional, Zero-Inflated, and Dispersion parameters. The Conditional parameters estimate the effect of fixed and random effects on the number of nestlings produced, the Zero-Inflated parameters estimate a zero-inflated intercept term to account for increased probability of producing zero nestlings relative to a standard Poisson distribution, and the Dispersion intercept estimates the deviation of the models from the expected Poisson-distribution dispersion (expected dispersion = 1, underdispersed < 1, overdispersed > 1). We present $\beta$ estimates for fixed effects with the standard error below in parentheses, standard deviations for random effects, sample sizes, and information on model fits.
